# Loss of heterozygosity spectrum varies with ploidy levels in natural yeast populations

**DOI:** 10.1101/2022.05.23.493037

**Authors:** Abhishek Dutta, Fabien Dutreux, Joseph Schacherer

## Abstract

The appearance of genomic variations such as loss of heterozygosity (LOH) has a significant impact on phenotypic diversity observed in a population. Recent large-scale yeast population genomic surveys have shown a high frequency of these events in natural isolates and more particularly in polyploids. However, the frequency, extent, and spectrum of LOH in polyploid organisms have never been explored and is poorly characterized to date. Here, we accumulated 5,163 LOH events over 1,875 generations in 76 mutation accumulation (MA) lines comprising nine natural heterozygous diploid, triploid, and tetraploid natural *S. cerevisiae* isolates from different ecological and geographical origins. We found that the rate and spectrum of LOH are variable across ploidy levels. Of the total accumulated LOH events, 8.5%, 21%, and 70.5% of them were found in diploid, triploid, and tetraploid MA lines, respectively. Our results clearly shows that the frequency of generated LOH events increases with ploidy level. In fact, the cumulative LOH rates were estimated to be 9.3 × 10^−3^, 2.2 × 10^−2^, and 8.4 × 10^−2^ events per division for diploids, triploids, and tetraploids, respectively. In addition, a clear bias towards the accumulation of interstitial and short LOH tracts is observed in triploids and tetraploids compared to diploids. The variation of the frequency and spectrum of LOH events across ploidy level could be related to the genomic instability, characterizing higher ploidy isolates.

**Highlights:**

- Yeast isolates with a high ploidy level (*e*.*g*., 3n and 4n) accumulate a larger number of LOH events compared to diploids.
- The type of events and the LOH spectrum are both variable according to the ploidy levels.
- A clear bias towards the accumulation of short LOH tracts is observed in polyploids compared to diploids.
- The variation in LOH spectrum is most likely related to genomic instability present in higher ploidy strains.

## Introduction

Error-prone DNA recombination and repair can impair fitness but might be counterbalanced by mechanisms that lead to loss of heterozygosity (LOH) (Korpanty et al., 2011; Lancaster et al., 2019; Moehring, 2011; Strathern et al., 1995; van Helden, 2012). In fact, LOH events can promote the expression of recessive alleles and generate novel allelic combinations, resulting in an increased genetic diversity and the emergence of new genotypes and therefore phenotypes. The mechanisms and evolutionary repercussions of LOH accumulation have been explored in several model organisms, primarily the *Saccharomyces cerevisiae* yeast (Dutta et al., 2021; James et al., 2019; Lee et al., 2009; Loeillet et al., 2020; St Charles and Petes, 2013; Sui et al., 2020). LOH events are categorized into interstitial, terminal, and whole chromosome-LOH. Interstitial or short-range LOH events are outcomes of mechanisms leading to gene conversions or double crossovers (Sui et al., 2020). Reciprocal crossovers (RCO) followed by co-segregation of identical alleles and break-induced replication (BIR) result in terminal or long-range LOH events (Sui et al., 2020). In contrast, in whole-chromosome LOH, one homolog is lost due chromosomal non-disjunction, alternatively, the remaining chromosome can be duplicated by re-replication. Whole-chromosome LOH is often a downstream consequence of aneuploidy (Chapman et al., 2012; Llorente et al., 2008; Pâques and Haber, 1999; Symington et al., 2014).

More recently, mutation accumulation (MA) and serial passaging experiments have been widely employed to elucidate the genome-wide dynamics of LOH accumulation and their phenotypic consequences in different intra-or inter-specific diploid yeast hybrids (Charron et al., 2019; Dutta et al., 2017; Ene et al., 2018; James et al., 2019; Loeillet et al., 2020; Nguyen et al., 2020; Smukowski Heil et al., 2017; Sui et al., 2020). Globally, LOH events, point mutations, and short indels (<100 bp) were the most frequently detected genetic alterations in yeast diploid genome when the selection is minimal during vegetative propagation (Loeillet et al., 2020; Sharp et al., 2018; Sui et al., 2020).And among them, the LOH event frequency has been estimated to be up to 6 orders of magnitude higher (Dutta et al., 2021). Further exploration even revealed diverse signatures or spectra in distinct genetic backgrounds, each defined by a different pattern of interstitial and terminal LOH events (Dutta et al., 2021; Loeillet et al., 2020; Nguyen et al., 2020). In addition, bursts of genomic instability leading to a large number of LOH events and a complete homozygotization have been highlighted during vegetative progression (Dutta et al., 2021; Sampaio et al., 2020).

While very fruitful, an important limitation of these studies has been the use of laboratory-generated artificial diploid hybrids with uniform levels of heterozygosity (Dutta et al., 2021, 2017; James et al., 2019; Loeillet et al., 2020; Pankajam et al., 2020; Sui et al., 2020). No survey has focused on LOH accumulation and its potential variability in natural isolates showing different ploidy levels. Natural *S. cerevisiae* isolates are mainly diploid, but polyploid isolates (3-5n) are frequent and enriched in specific subpopulations such as the beer clade (Gallone et al., 2016; Saada et al., 2022). In addition, other yeast species (such as *C. neoformans* and *C. albicans*) are predominantly polyploids and LOH events are widespread (up to 20% of the genome) (Abbey et al., 2014; Feldmesser et al., 2000; Mba et al., 2022; Selmecki et al., 2010; Smith and Hickman, 2020).

Here, we therefore conducted a MA experiment to have a better idea of the genome-wide LOH landscape in natural diploid and polyploid *S. cerevisiae* isolates, coming from various ecological and geographical origins. MA lines derived from diploids, triploids, and tetraploids were evolved through 75 single-cell bottlenecks (1,875 generations). Our analysis revealed that the LOH rate and spectrum highly depend on the ploidy level. LOH events occur more frequently in isolates with higher ploidy, with a LOH rate of 9.3 × 10^−3^, 2.2 × 10^−2^, and 8.4 × 10^−2^ events per division for diploids, triploids, and tetraploids, respectively. In fact, polyploids accumulate more internal and shorter LOH tracts.

## Results

### Experimental design of the natural MA lines

Using *S. cerevisiae* as an experimental model, we characterized the genome-wide landscape and dynamics of LOH events in natural diploids as well as polyploid isolates. A total of 76 MA lines were analyzed covering nine different heterozygous genetic backgrounds with three ploidy levels: diploid (n=25), triploid (n=24), and tetraploid (n=27) (Table 1; Figure S1). The natural *S. cerevisiae* isolates were chosen from various ecological (*e*.*g*., beer, wine, and clinical) and geographical origins (*e*.*g*., Africa, South America, and Europe), and designated as ancestors (Table S1) (Peter et al., 2018). Replicate lines were generated from each of the ancestral isolates to set up an MA experiment. The MA lines were propagated purely vegetatively and subjected to single-cell bottlenecks every 48 hours for 75 bottlenecks on rich media (YPD). We estimated the MA lines to have undergone an average of approximately 25 divisions per bottleneck for a total of at least 1,875 divisions per line (Dutta et al., 2021; Liu and Zhang, 2019). Selection is minimal and genomic changes have accumulated solely due to drift. At the end of the experiment, the genomes of the 76 lines were completely sequenced using a short-read Illumina strategy.

**Table 1:** Natural MA lines. * Standardized names from Peter et al 2018

| **Ploidy** | **Std.**<br>**Name\*** | **Het**<br>**positions** | **No. of**<br>**sequenced**<br>**lines** | **No. of**<br>**bottlenecks** |
| --- | --- | --- | --- | --- |
| **Diploid** | BDL | 39220 | 08 | 75 |
| **Diploid** | BIE | 55580 | 11 | 75 |
| **Diploid** | CBL | 56485 | 07 | 75 |
| **Triploid** | BEN | 19549 | 13 | 75 |
| **Triploid** | ARN | 39985 | 06 | 75 |
| **Triploid** | ATB | 60269 | 05 | 75 |
| **Tetraploid** | BEK | 26889 | 09 | 75 |
| **Tetraploid** | CBR | 53143 | 08 | 75 |
| **Tetraploid** | CKK | 77268 | 11 | 75 |

### Variable and ploidy-dependent LOH spectra in *S. cerevisiae*

We first identified and determined the LOH events present in the 76 MA lines. LOH events were defined as regions with a length of more than 900 bp and at least three consecutive converted Single Nucleotide Polymorphisms (SNPs) (see Materials and methods). A 900 bp threshold was defined to overcome biases due to the variable level of heterozygosities (Figure S3). Events greater than 900 bp and supported by at least three consecutive converted SNPs were defined as being under LOH (see Materials and methods; Figure S3). Adjacent LOH events were merged if at least two consecutive heterozygous SNPs did not support the disruption. The size of a LOH event corresponds to the distance separating the midpoint between the closest upstream unconverted and converted sites and the midpoint between the closest downstream converted and unconverted sites. From the MA lines data, we compiled a genome-wide LOH map across the three ploidy groups by masking the homozygous regions present in the ancestral isolates (Figure 1).

**Figure 1.**
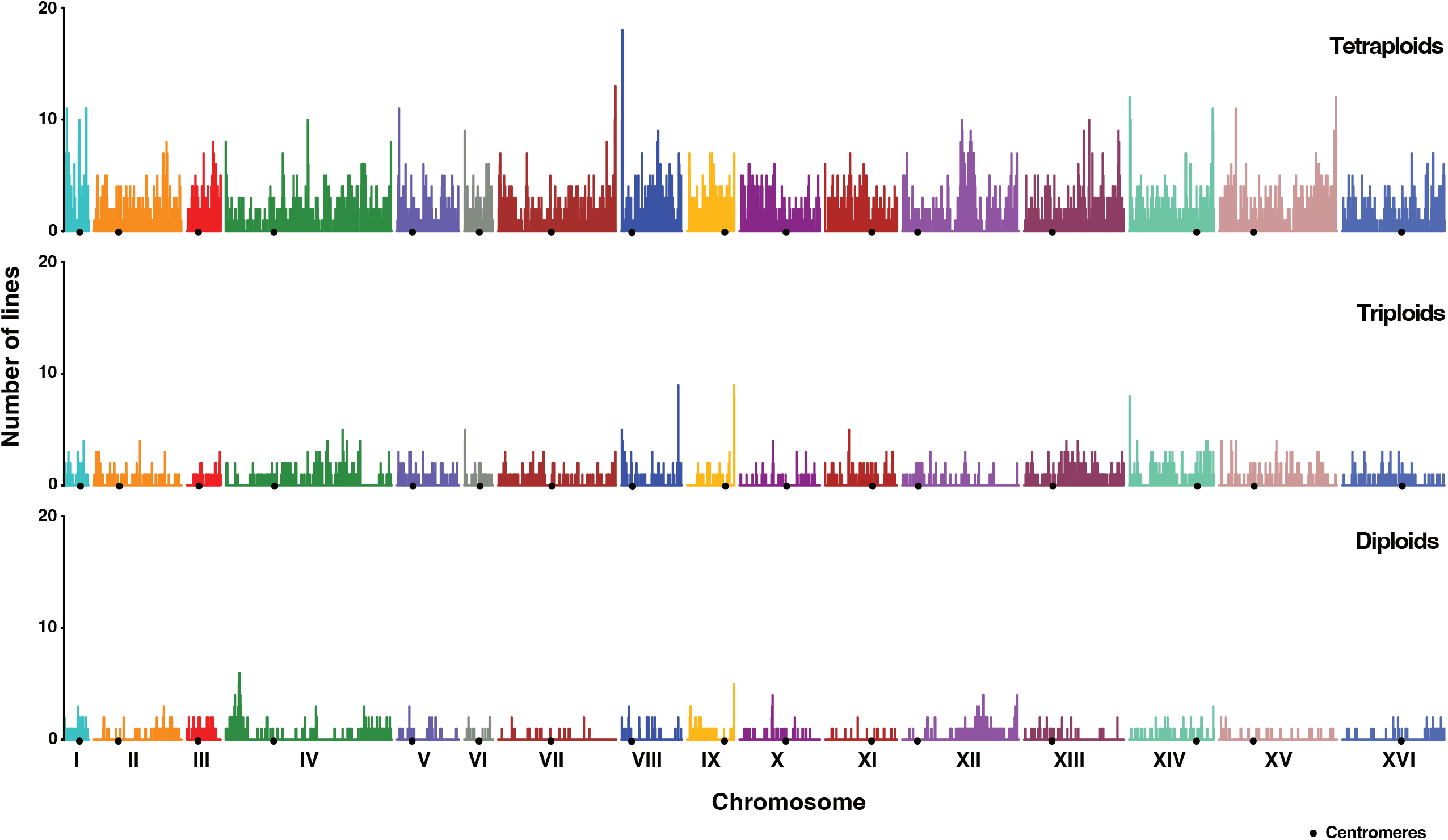
Chromosome-wide map representing cumulative frequencies of regions under LOH in the diploid, triploid, and tetraploid MA lines.

A total of 5,163 events were identified in the 76 MA lines, 437, 1,087 and 3,639 of which were found in the diploid, triploid, and tetraploid MA lines, respectively (Table S2; Figures 2A; S4A). As previously observed for the artificial laboratory diploids (Dutta et al., 2021), interstitial events were significantly more frequent than terminal events regardless of ploidy level (Figure S4A). However, the total number of the two types (interstitial and terminal) per line significantly increase with the ploidy (Figure 2A; Table S2). These data also allowed us to estimate the LOH rates and we found a rate of 9.3 × 10^−3^, 2.2 × 10^−2^, and 8.4 × 10^−2^ events per division for the diploids, triploids, and tetraploids, respectively. By determining the rates of interstitial and terminal LOH events, we found the same trend with a ploidy level dependent increase.

**Figure 2.**
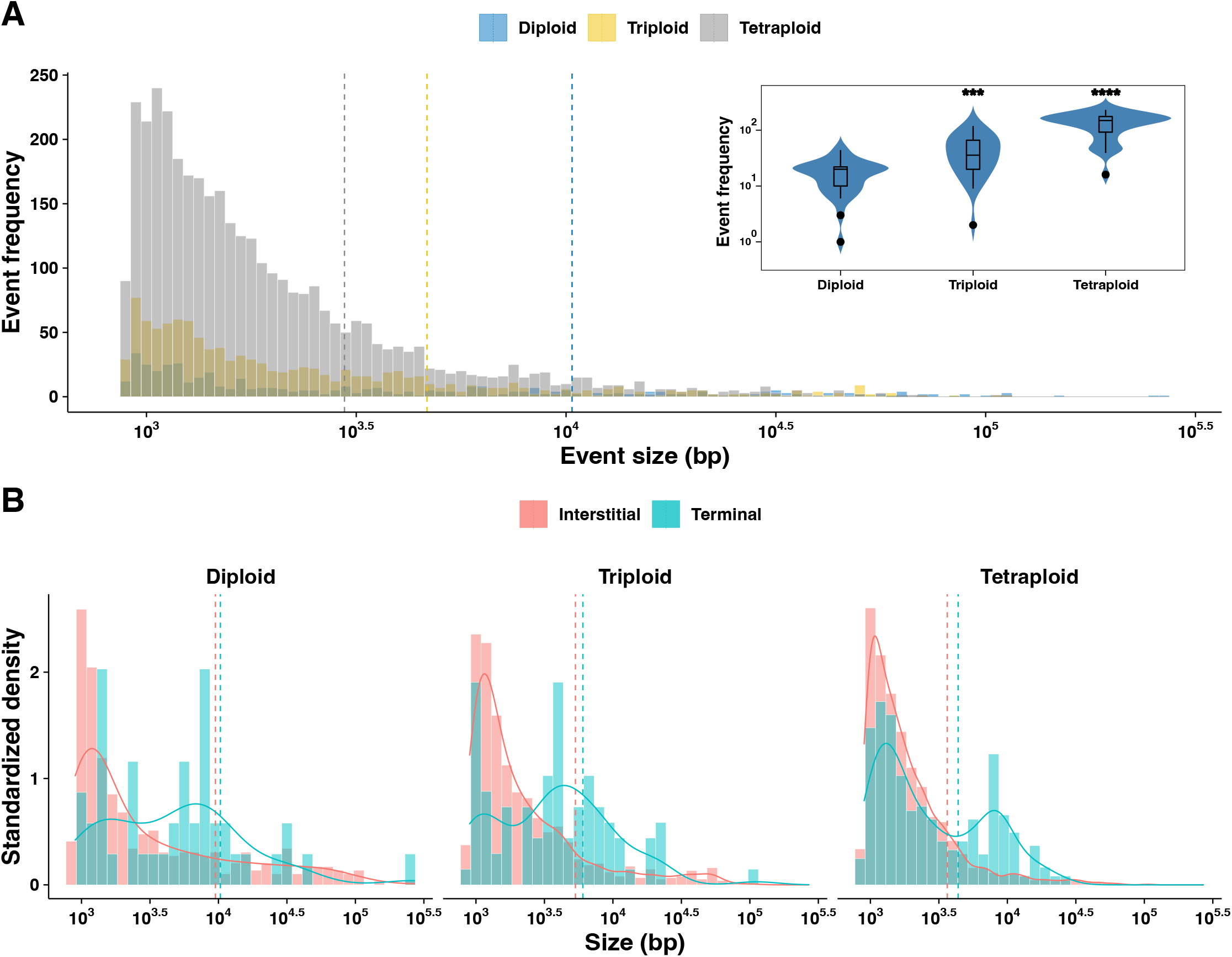
Ploidy dependent variability in LOH accumulation. **A**. Overall ploidy wise size distribution of the LOH tracts in the 76 MA lines. The overall average LOH event sizes were **10.3 kb, 4.6 kb** and **2.9 kb** respectively for the diploids, triploids, and the tetraploids. **[Inset:** Violin plot of the LOH event counts in the MA lines, triploid and tetraploid lines accumulate significantly greater LOH compared to the diploids (Wilcoxon test, ***p<0.001; ****-p<0.0001; ns-not significant).**] B**. Ploidy wise size distribution of the terminal LOH events and not the interstitial events was bimodal. The average tract lengths were **9.9 kb, 4.5 kb, 2.9 kb** and **13.3 kb, 6.6 kb, 4.3 kb** respectively for the interstitial and terminal events.

Subsequently, we looked at the size distribution of the LOH events. Overall, the size of the LOH events was 10.3 kb, 4.7 kb, and 2.9 kb on average for the diploid, triploid, and tetraploid MA lines, respectively. In fact, we observed a ploidy-dependent decrease in LOH event sizes (Figure 2A-B; S4B; Table S2). As expected, terminal events were always larger than interstitial events regardless ploidy level (Figure S4B). Event size variance was consistently high, but highest in the diploids (CV: 245%; Levene’s Test, p<2.2 × 10^−16^), followed by the triploids (CV: 213%; Levene’s Test, p<2.2 × 10^−16^) and tetraploids (CV: 182%; Levene’s Test, p<2.2 × 10^−16^). A chromosome size-dependent increase in LOH events was observed for all ploidy levels but only for interstitial events (Figure S5).

Based on the number and size of the events, we estimated the accumulated fraction of the genome under LOH. The acquired genome under LOH was 2.3% on average and ranged from 0.01-8%. The polyploid MA lines displayed a larger fraction of genome under LOH (2.7% on average) compared to the diploid lines (1.5% on average) (Figure S6; Wilcoxon test, p<0.01). Nevertheless, the acquired fraction of genome under LOH was much greater (10.5% on average) in the artificial lines we generated in a previous survey (Dutta et al. 2021). Although slightly different, it clearly demonstrates that the accumulated LOH faction is of the same order of magnitude in natural polyploid and diploid isolates.

Overall, the LOH event frequency (interstitial, terminal, and total) was strongly correlated with ploidy level (Spearman’s R=0.8; p<2.2 × 10^−16^). Only the interstitial event size negatively correlated with ploidy (Spearman’s R=-0.4; p<0.001) while the terminal event sizes were unaffected (Spearman’s R=-0.1; p>0.05) (Figure S5). Moreover, the ratio between interstitial and terminal events is similar whatever the level of ploidy (Figure S4C; Table S2) (p> 0.05; Kruskal-Wallis test).

### Natural hybrids accumulate less LOH events than artificial hybrids

We recently explored the accumulation of LOH events in artificial diploid *S. cerevisiae* hybrids generated via crosses between stable haploids, representing different genetic backgrounds (Dutta et al. 2021). We therefore compared the variation of the LOH spectra obtained for natural heterozygous diploid isolates and these artificially generated hybrids. For that purpose, we compared the LOH landscape of three artificial hybrids (BTI/ABA, ACD/AKQ and ACK/CMQ) having a similar heterozygosity level to that of the natural diploids studied here.

By comparing these set of MA lines, we found that the size distribution of the LOH events was similar for the artificial and natural lines (Figure 3A). The average size of LOH events (including interstitial and terminal events) was 11.1 kb and 10.3 kb for artificial and natural MA lines respectively, clearly showing no difference (Figure 3C). However, the artificial diploid lines had a significantly higher frequency of events (65.9 events per line on average) than natural diploids (16.8 events per line on average) (Wilcoxon test; p<2 × 10^−16^) (Figure 3B). In fact, the LOH rate was 3.2 × 10^−2^ events per division and 9 × 10^−3^ events per division for the artificial and natural lines, respctively. Moreover, both interstitial and terminal event rates were at least 3-fold higher in the artificial lines.

**Figure 3.**
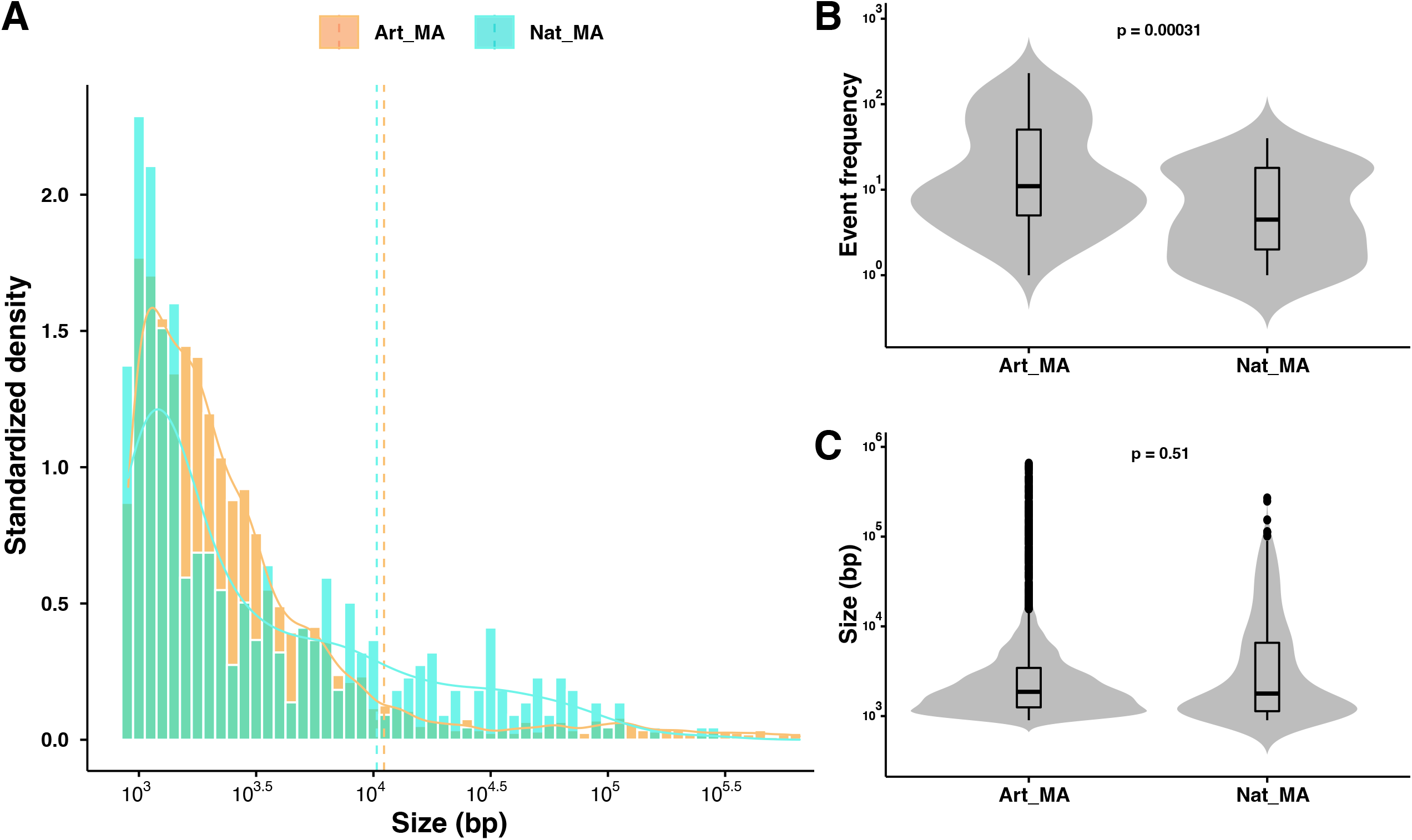
LOH accumulation patterns in artificial vs natural diploids within similar heterozygosity levels. **A**. The size distributions trends are similar in MA lines derived from artificial (average size-**11.4 kb**) and natural diploids (average size-**10.3 kb**). **B**. Violin plot of the LOH event counts in artificial vs natural diploid MA lines. Artificial diploid MA lines accumulate significantly greater interstitial and terminal LOH events (Wilcoxon test, *-p<0.05; **-p<0.01; ***p<0.001; ****-p<0.0001; ns-not significant). **C**. Violin plot of the LOH event sizes in artificial vs natural diploid MA lines. Interstitial and terminal LOH event sizes are similar (Wilcoxon test, *-p<0.05; **-p<0.01; ***p<0.001; ****-p<0.0001; ns-not significant).

Similarly, the acquired fraction of genome under LOH was far greater in the artificial lines (10.5%) compared to the natural lines (1.5%) (Wilcoxon test, p<0.01). Interestingly, no nearly homozygous lines (> 90 % of the genome under LOH) were found in the natural lines, while 5% of the artificial lines did, probably highlighting a higher degree of instability in the artificial hybrid genomes.

### Chromosomal instabilities in the MA lines

New generated heterozygous and polyploid hybrid genomes generally show a higher rate of genetic instability (Gerstein et al., 2008; Morales and Dujon, 2012). Therefore, polyploidy recurrently leads to aneuploidy and genetic diversity, and has been considered a major evolutionary driver. However, the rate of genetic instability in natural diploid and polyploid isolates is not well characterized.Artificially generated diploid and polyploid hybrids have shown to increase or decrease their overall ploidy level during propagation (Charron et al., 2019; Marsit et al., 2021). Moreover, experimentally generated tetraploid *S. cerevisiae* strains frequently revert to a diploid state through loss of chromosomes during mitotic progression (Gerstein et al., 2008; Selmecki et al., 2015). Here, we assessed the ploidy of all lines using flow cytometry, we found no substantial deviations from the ancestral ploidy level in the MA lines at the end of the experiment. This observation clearly indicates a stability of the ploidy level in the natural isolates over a mitotic propagation. It indicates that either the recurrent bottlenecks in the MA experiment prohibit ploidy changes from being selected, or that certain genetic backgrounds are not susceptible to spontaneous ploidy variations (Sharp et al., 2018). We then used sequencing coverage across the genome in the MA lines to detect changes in copy number variation, more precisely aneuploidies, resulting from mitotic nondisjunction. In total, we detected 83 chromosomal aneuploidy events in our 76 MA lines (Figure 4A-B). Aneuploidies were detected in 36 % (n=12) and 90% (n=71) of the diploid and polyploid lines, respectively (Table S3). The incidence of aneuploidies was found to be associated with ploidy in the MA experiment (Spearman’s R= 0.3; p<2.2 × 10^−16^). The rates of aneuploidies were 2.6 × 10^−4^ events per division and 7.4 × 10^−3^ events per division in the diploids and polyploids respectively, highlighting a 28-fold difference. All chromosomes were involved in aneuploidy events, with the exception of chromosome XVI. Additionally, we observed no difference in the LOH event rates in the euploid and aneuploid lines (p>0.05, Wilcoxon test).

**Figure 4.**
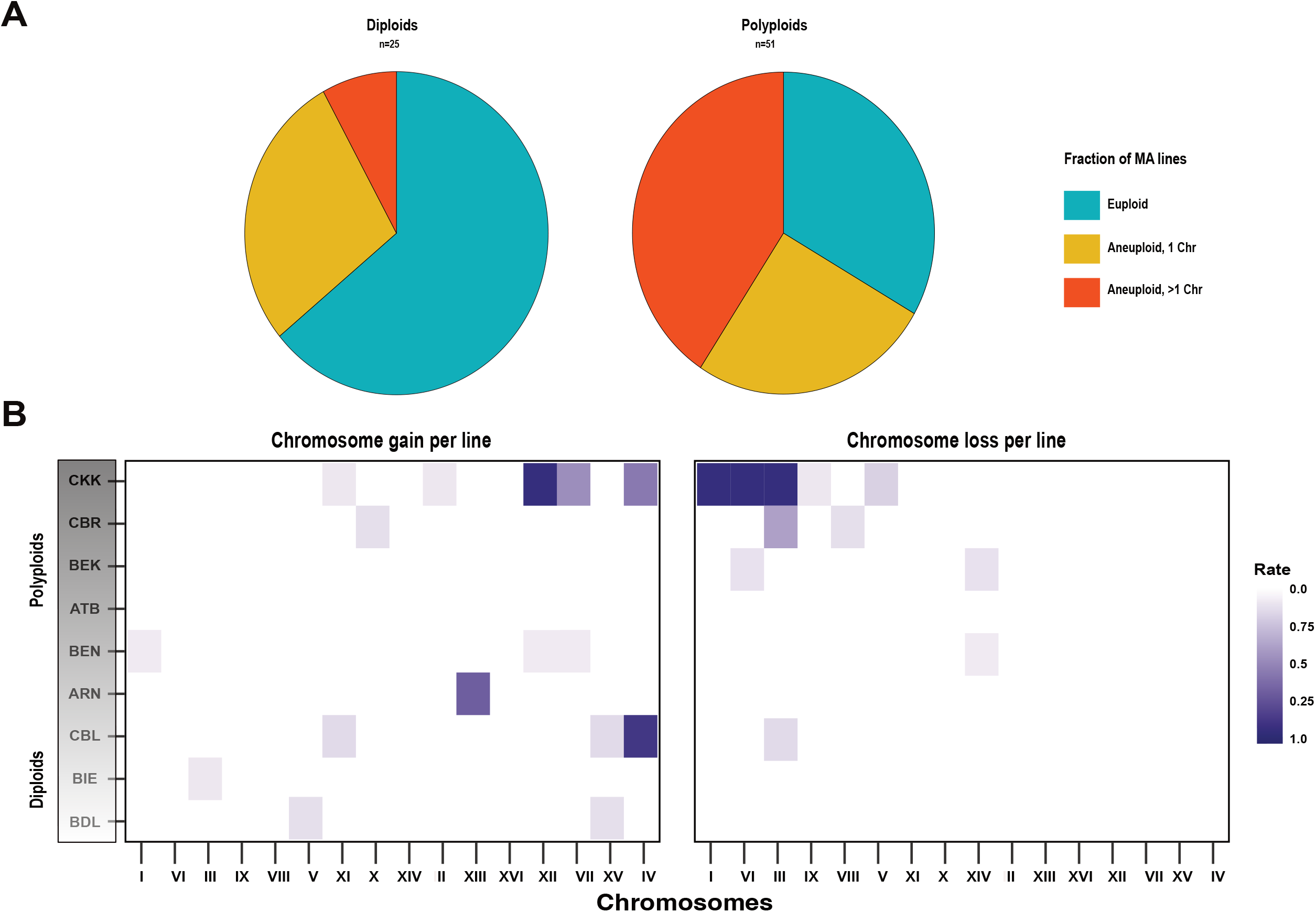
**A**. Pie chart representing fraction of MA lines euploid, aneuploid (+/-1 chromosome) or aneuploid (+/->1 chromosome). **B**. Frequency of aneuploidies per line across the individual backgrounds. **ATB** (3n) derived MA lines and Chromosome XVI overall was never involved in an aneuploidy event and have been excluded from the plot.

### Growth fitness and fertility of the MA lines are stable throughout propagation

We measured the evolution of mitotic growth and fertility of the MA lines over the course of the experiment *i*.*e*., bottlenecks, 25, 50 and 75. Our results show that there are both systematic gain and loss of fitness for the independent diploid lines (Table S4; Figure 5A). Interestingly, the polyploid MA lines never displayed any gain of fitness (Figure 5A; r= -0.2, p<0.05). Globally, we can observe a general gain of fitness in the diploid lines (R=0.2, p<0.01), while a decline in fitness of the polyploid lines (Figure 5A-B). Further, the change in fitness was neither significantly associated with LOH nor the aneuploidies.

**Figure 5.**
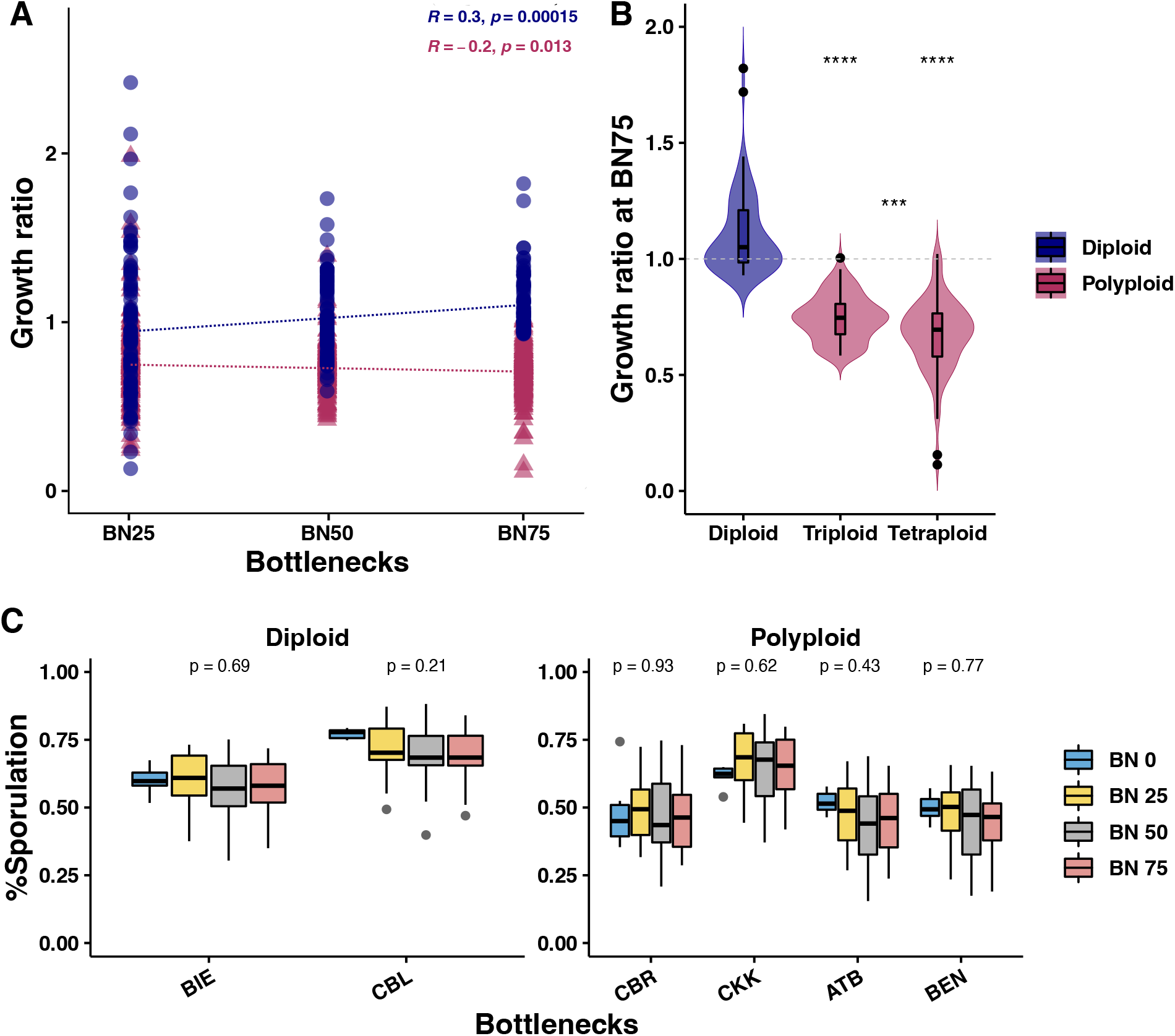
**A**. Mean growth ratio in the individual MA lines (bottlenecks 25, 50 and 75) compared to the ancestral isolates (bottleneck 0) traced over the course of the experiment. Diploid MA lines display positive growth trend (Spearman’s, R= 0.3; p<0.01), while polyploids display a negative growth trend (Spearman’s, R= -0.2; p=0.013). **B**. Violin plot representing growth in the MA lines at the end of the experiment (Wilcoxon test, *-p<0.05; **-p<0.01; ***p<0.001; ****-p<0.0001; ns-not significant). **C**. Sporulation efficiencies do not change in the evolved MA lines compared to the ancestral isolates. BDL, ARN and BEK ancestral isolates as well as the evolved MA lines did not sporulate. Spore viability percentages at bottlenecks 0 and 75 shown.

We then compared the evolution of fertility during the MA lines propagation. We measured both the sporulation efficiency and spore viability components of fertility in these lines. In diploids, meiotic spore fertility negatively correlates with divergence and the gradual restoration can be facilitated by accumulating LOH events (Cubillos et al., 2011; Dutta et al., 2021, 2017). In contrast, evolved tetraploids post whole genome duplication (WGD) ensured fertility rescue in highly diverged inter-specific *S. cerevisiae* / *S. paradoxus* hybrids (Charron et al. 2019; Marsit et al. 2021). Moreover, in a previous study, we found significant fertility recovery to be associated with accumulation of short interstitial LOH events, as observed in the triploid and polyploid MA lines (Dutta et al., 2021). In the ancestral backgrounds that could sporulate, we tested the sporulation ability of all the derived lines and traced the evolution of the sporulation efficiency for three MA lines, chosen at random. MA lines retained their ability to sporulate and in fact, we found no significant differences in the evolved lines at the different bottlenecks (Table S5; Figure 5C). Next, we measured the meiotic spore fertility of 18 lines across the different genetic backgrounds and ploidy groups at the end of the experiment. Surprisingly, we observed no significant improvements in either the diploid, or polyploid MA lines (Figure 5C).

## Discussion

The patterns of LOH accumulation in wild *S. cerevisiae* isolates have never been explored and most insights in this regard come from laboratory generated inter-or intra-specific hybrids. The scale and role of LOH events in hybrid genome stabilization is not well understood in a natural diploid as well as polyploid context. In fact, *S. cerevisiae* and *C. albicans* polyploid isolates have been isolated from diverse ecological and geographical origins, suggesting that they alter their ploidy states to adapt to changing environments (Peter et al., 2018; Todd et al., 2019). Laboratory generated polyploids display rapid ploidy reductions and chromosomal instabilities in evolution experiments (Gerstein et al., 2008; Selmecki et al., 2015). The persistence of polyploids and their evolution is therefore intriguing. Here, we explored the LOH spectrum and chromosomal instabilities in natural heterozygous diploid and polyploid isolates through a MA experiment. Our results demonstrate that polyploids accumulate more LOH events compared to diploids after 1,875 generations. In addition, polyploid MA lines mainly accumulate very short LOH events. Furthermore, we observed that the artificial hybrids exhibit a higher LOH event rate compared to natural isolates. However, the size and distribution of these events are similar to those found in the natural diploid MA lines (Dutta et al., 2021; Sui et al., 2020). Overall, the ploidy level of MA lines generated from wild strains is extremely stable, according to our results. MA polyploid lines showed strong chromosomal instability but the fitness and fertility of these MA lines did not change and is not correlated with LOH or chromosomal instabilities.

Polyploidy itself has been associated with increased genomic instability in yeast (Storchova, 2014; Selmecki et al., 2015; Marsit et al., 2021). In fact, polyploid cells in *Drosophila melanogaster* have been shown to display an elevated DNA damage response (Nandakumar et al., 2020). Tetraploid genomes have been shown to be more sensitive to Double Strand Breaks (DSBs) than diploids and furthermore, heterozygosity is known to hamper the DSB repair process (Skoneczna et al., 2015; Strathern et al., 1995). The high rates of LOH events in polyploid lines may be explained by the fact that heterozygous polyploid genomes may be associated with high DNA damage, which upon recombination repair results in increased LOH events.

A striking observation is the decrease in size of LOH events with increasing ploidy level. The size of these events were 10.3 kb and 4.3 kb in diploid and polyploid lines, respectively. Similarly, artificial *S. cerevisiae* / *S. paradoxus* hybrid tetraploids that spontaneously evolved from ancestral hybrid diploids in an MA experiment displayed shorter events compared to their diploid counterparts (Marsit et al., 2021). Utilizing a microarray-based approach, Yim et al., (2014) described two distinct classes of conversion events with median sizes of 6 kb (more frequent) and 54 kb (less frequent), arising from different DNA lesions and repair mechanisms. Prior studies have also shown heteroduplex formation, followed by mismatch repair, results in short mitotic conversion events less than 1 kb (Mitchel et al., 2010). The high frequency of LOH events suggests that polyploids undergo more breaks.

Moreover, given the sensitivity of yeast polyploids to DSBs, the involvement of long stretches of DNA in conversions can be deleterious to pairing and disentangling and disrupt ancestral linkage groups. In addition, the size of conversion events is under strong regulation of the MutLβ complex as proposed during meiotic recombination (Duroc et al., 2017). Finally, mutants defective in MMR activity as well as overexpression of MMR proteins have also been proposed to impact the size of conversion events (Campbell et al., 2014; Oke et al., 2014).

We also found that chromosomal instability is higher in polyploids compared to diploids. The rates of aneuploidy were within range of previous estimates in *S. cerevisiae* diploids, but significantly lower compared to laboratory generated tetraploids (Gilchrist and Stelkens, 2019; Sui et al., 2020; Marsit et al., 2021; Dutta et al., 2021). A recent study has also suggested very high genomic stability and a relatively low incidence of aneuploidies in triploid and tetraploid industrial *S. cerevisiae* isolates post return to growth (RTG) (Mozzachiodi et al., 2021). Increased incidence of aneuploidies compared to diploids may also be a consequence of recombination and multivalent segregation. Interestingly, the patterns of aneuploidies were different in polyploids compared to diploids. Polyploids show more aneuploidies of medium and large chromosomes, and the frequency of chromosome loss is significantly higher. By contrast, we observed aneuploidies to be associated with the small chromosomes and losses to be rare in diploids. This suggests that diploid isolates are more impacted by chromosomal imbalances.

Finally, LOH accumulation and chromosomal instabilities had no effect on the mitotic growth and fertility of the evolved lines. Overall, our results suggests that in natural polyploids, the ploidy level does not vary, probably because the genomic states have been selected for and evolved over a long course of time. In polyploids, LOH accumulation is more important than in natural diploids leading to a higher genetic diversity. Nevertheless, the phenotypic consequences of this accumulation still need to be assessed.

## Materials and methods

### Strain construction and MA lines propagation

Tables 1 and S1 summarize the *S. cerevisiae* strains utilized in this study. Yeast strains were grown on either YPD (yeast extract 1%, peptone 2%, dextrose 2 %) or synthetic complete (yeast nitrogen base 0.67%, amino acid mix 0.2%, dextrose 2%) medium at 30°C. Single colonies were isolated from each of the ancestral isolates, and ploidies were confirmed by flow cytometry. These were also checked for their ability to sporulate on 1% potassium acetate-agar. Individual replicate lines were bottlenecked to a single colony every 48 hours for at 75 bottlenecks on YPD agar. Intermediates were frozen down every 25 bottlenecks until the end of the experiment. MA lines at the end of the experiment were sequenced. The ploidies of the MA lines were verified via flow cytometry (BD Accuri C6, BD biosciences) before and at end of the MA lines experiment to verify the ploidy level.

### Determining sporulation efficiencies and growth

All MA lines and ancestral isolates were streaked down to single colonies on YPD agar plates. On sporulation media (1% potassium acetate agar), three separate colonies from each of the lines and ancestral isolates were patched (Argueso et al., 2004). To evaluate sporulation efficiency, 100 cells per sample were counted using an optical microscope at the end of 72 hours. The number of dyads, triads, and tetrads divided by the total cell count was used to calculate sporulation efficiency. Mitotic growth was determined using endpoint colony growth on solid media. MA lines were pre-cultured in YPD broth and pinned onto a solid YPD plate to a 1536 density format using the replicating ROTOR robot (Singer Instruments). The plates were incubated for 24 hours at 30°C and were scanned with a resolution of 600 dpi at 16-bit grayscale. Quantification of the colony size was performed using the R package Gitter and the growth of each of the lines was measured by estimating the growth ratio between the colony size of the MA line at bottlenecks 25, 50, 75 to the respective colony size of the MA line at bottleneck 0 (Wagih and Parts, 2014).

### Whole-genome sequencing of MA lines

Genomic DNA was extracted from the MA lines using the Omega yeast DNA kit (Life Science Products). DNA libraries were prepared from 5 ng of total genomic DNA using the NEBNext Ultra II FS DNA Library Kit for Illumina (New England Biolabs) following manufacturers’ protocols. Following quality check using a Bioanalyzer 2100 (Agilent Technologies) and quantification using the Qubit dsDNA HS assay, 4 nM of each library were pooled and run on a NextSeq 500 sequencer with paired-end 150 bp reads by the EMBL Genomics Core Facility (Heidelberg, Germany).

### Read mapping, genotyping of sequencing data

Sequencing reads from Fastq files were mapped to the masked (RepeatMasker, default parameters, masking simple repeats and low complexity regions) *S. cerevisiae* R64 reference genome using bwa mem (v0.7.17). The resulting bam files were sorted and indexed using SAMtools (v1.9). Duplicated reads were marked, and sample names were assigned using Picard (v2.18.14). Finally, GATK (v3.7.0) was used to realign remaining reads. Candidate variants were then called using GATK UnifiedGenotyper. The calling was done simultaneously for lines from the same background.

### Analysis of LOH tracts

After variant calling, SNPs called in each hybrid parental couple were first filtered (bcftools v1.9) to define a set of confident markers expected to be heterozygous in the MA lines. Positions with a single alternate allele, supported by at least 50 sequencing reads across alleles were kept as parental markers. Joint SNP calling was performed background wise as described previously. Bcftools isec was used to extract SNPs located at parental markers positions in all samples. In case of homozygosity at markers position, a parental origin tag was added by comparing the allele to parental alleles. Positions with GQ<20 was filtered out. Loss of heterozygosity events were called using an in-house script. Heterozygous and LOH tracts were initially defined by uninterrupted tracts of successive marker positions with the right tag (heterozygous, homozygous). Single-marker tracts were filtered out and subsequent tracts from the same origin were merged. Any tracts with 80% or more overlap and shared by at least 50% of the lines from the same ancestor were excluded. Average LOH tract coordinates were determined as the mean between the coordinates of first or last marker of a given tract and the first previous or next marker around that tract. LOH tracts were tagged as terminal if they overlapped the first or last 20kb of a chromosome and tagged as interstitial otherwise. Subsequently, regions under LOH were defined as events >900 bp supported by at least there converted SNPs. The 900 bp threshold was selected by determining, an overlap between the frequencies and sizes of events in the Natural MA lines in this study with the artificial diploid MA lines from Dutta et al. 2021, to overcome biases in event detection because of the varying levels of heterozygosities (Figure S3). LOH event rates per line were calculated as N/D, where N= number of events (Interstitial, Terminal, or total events) and D=total number of divisions/ bottlenecks which we assumed to be 25, based on previous literature.

## Data availability

All sequenced MA lines listed in Table S6 are available upon request. In addition, sequence data are available from National Centre for Biotechnology Information Sequence Read Archive under accession number: PRJEB45355.

## Acknowledgements

We are thankful to Bertrand Llorente for critical reading of the manuscript and thoughtful suggestions. This work was supported by the European Research Council (ERC Consolidator Grant 772505). J.S. is a Fellow of the University of Strasbourg Institute for Advanced Study (USIAS) and a member of the Institut Universitaire de France.

## Supplementary figures

**Figure S1**. Cartoon representing the experimental design.

**Figure S2**. Distribution of the heterozygous single nucleotide polymorphism (SNP) densities as a fraction of total heterozygous SNPs in 5 kb windows across the ancestral isolates as described in Table 1.

**Figure S3**. Number events across size thresholds in the natural MA lines (Diploid and polyploid) and diploid hybrid MA lines (Dutta et al. 2021).

**Figure S4. A**. Interstitial events are significantly more frequent than terminal events across all ploidy groups (Wilcoxon test, *-p<0.05; **-p<0.01; ***p<0.001; ****-p<0.0001; ns-not significant). **B**. Interstitial events are significantly shorter than terminal events across all ploidy groups (Wilcoxon test, *-p<0.05; **-p<0.01; ***p<0.001; ****-p<0.0001; ns-not significant). **C**. Overall fraction of interstitial and terminal events are similar across all ploidies (Wilcoxon test, *-p<0.05; **-p<0.01; ***p<0.001; ****-p<0.0001; ns-not significant).

**Figure S5**. The frequency of interstitial LOH events increase with increasing heterozygosity for diploids, triploids, and tetraploids.

**Figure S6**. The fraction of acquired genome under LOH in diploids and polyploids (Wilcoxon test, *-p<0.05; **-p<0.01; ***p<0.001; ****-p<0.0001; ns-not significant).

