## Supplementary figures and images for "Loss of heterozygosity spectrum varies with ploidy levels in natural yeast populations"

### Figure S1

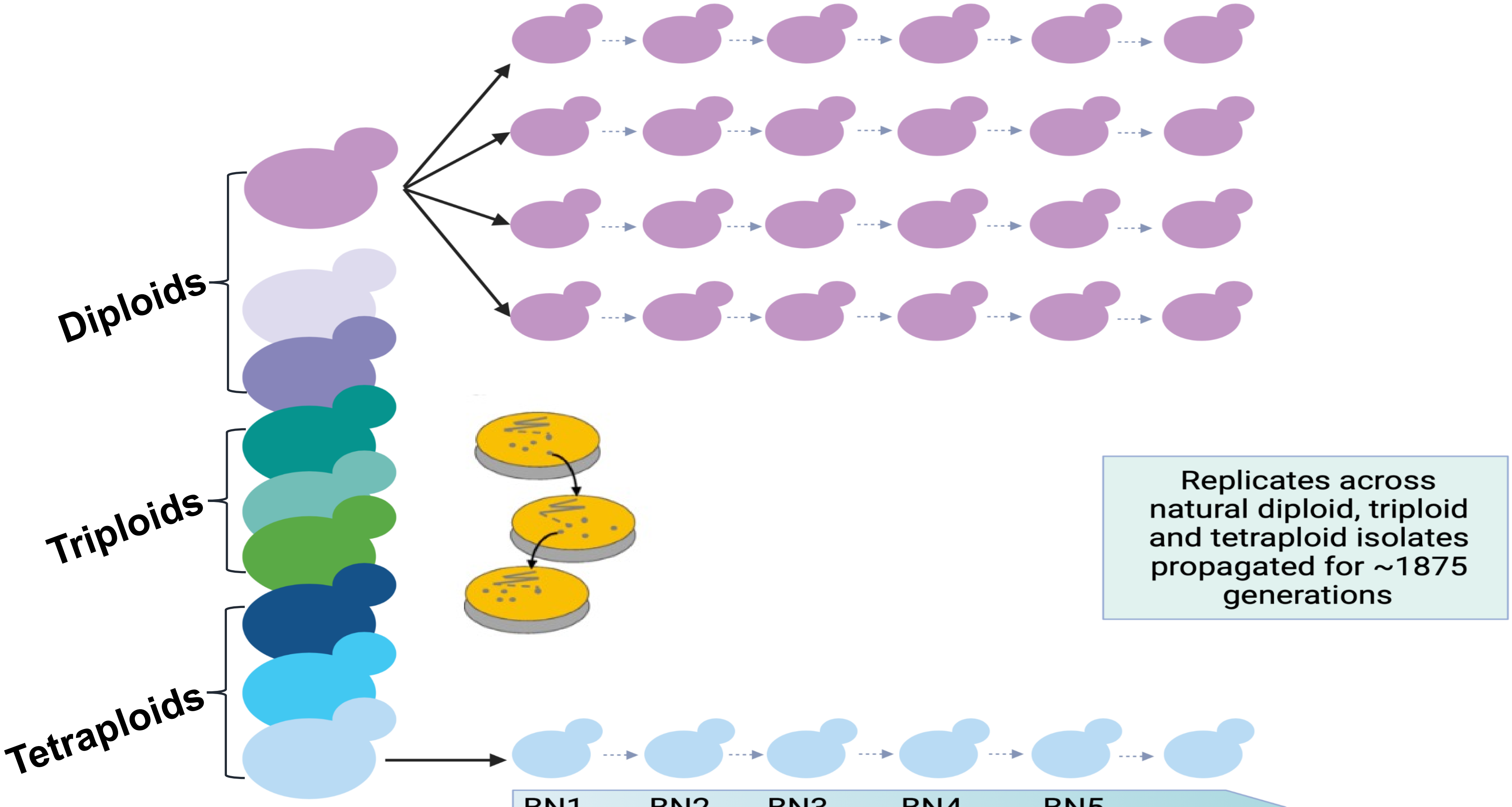

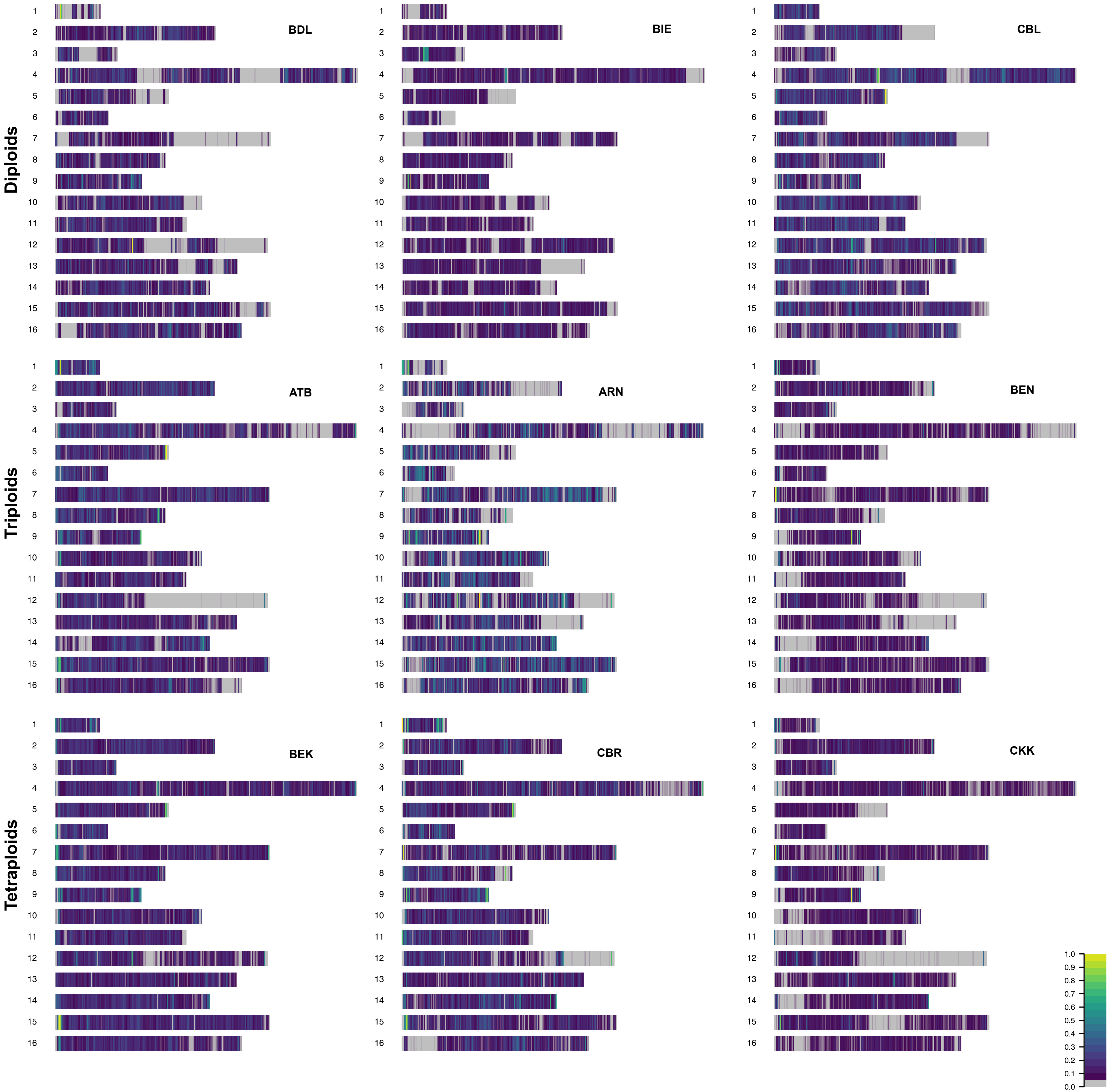

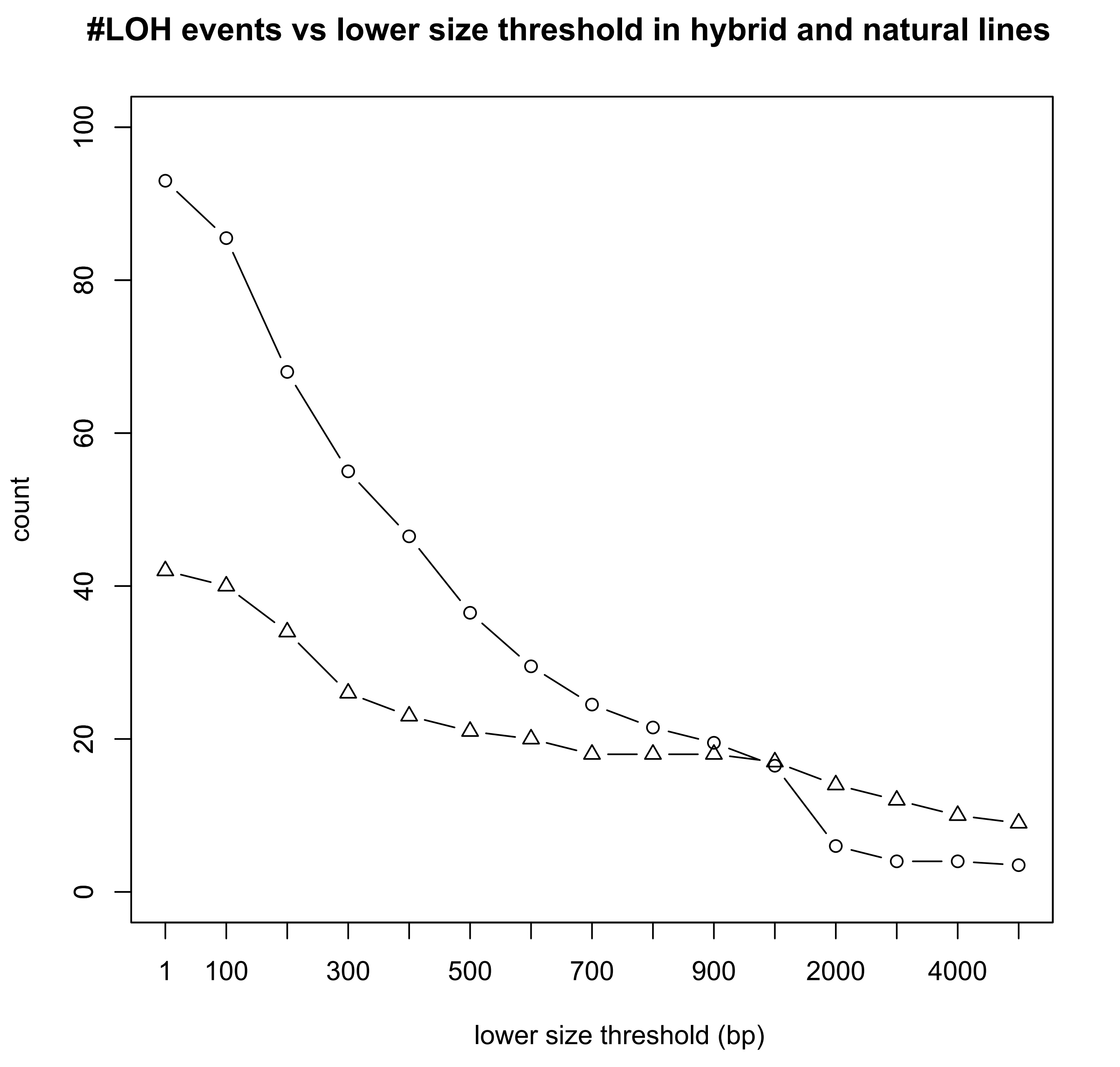

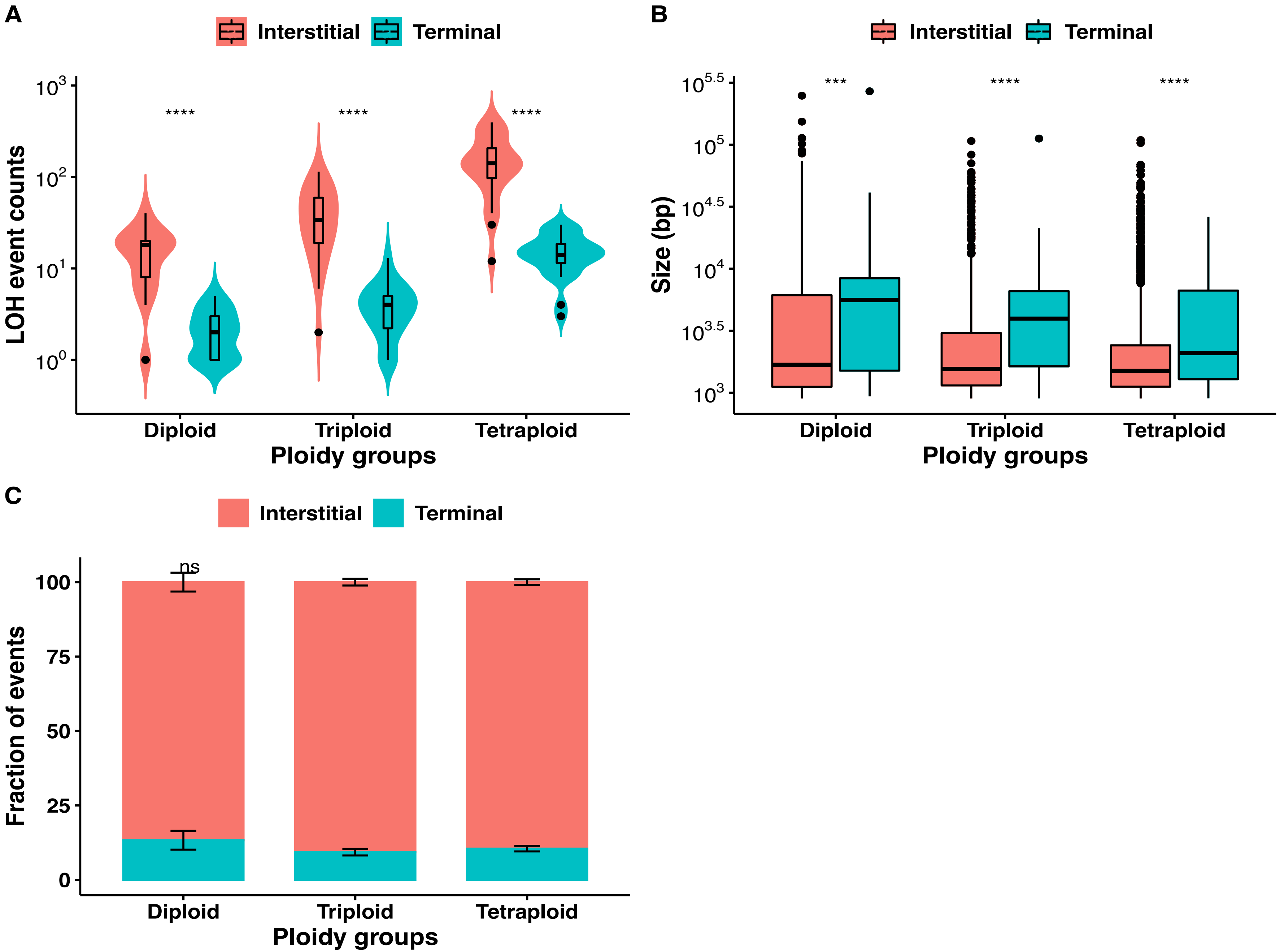

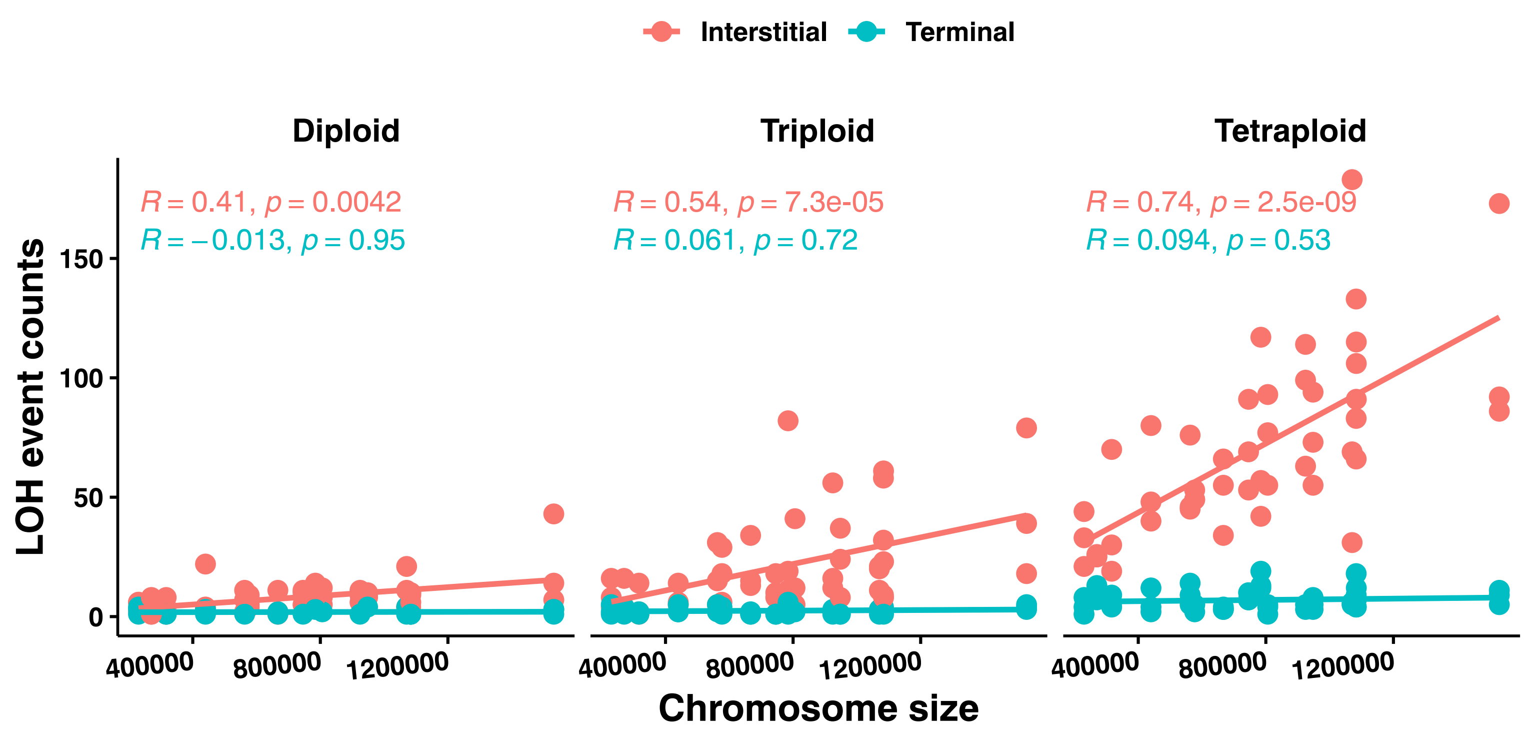

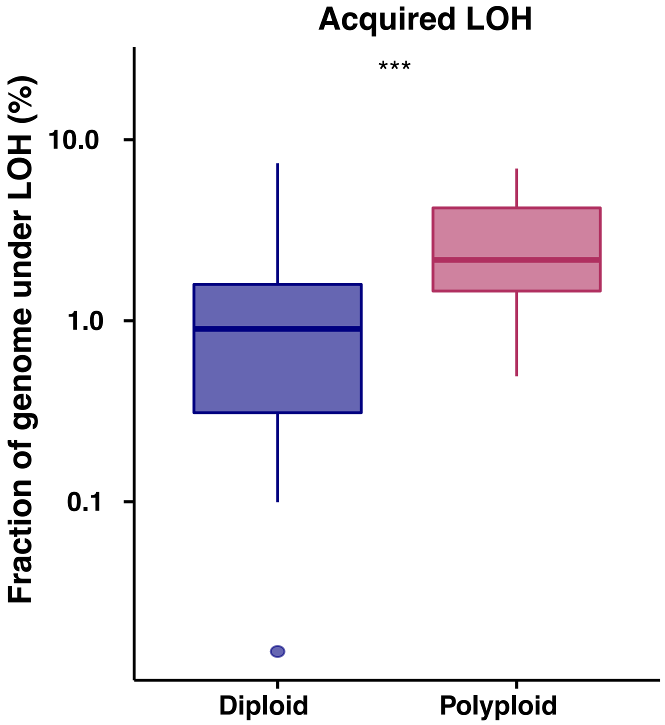
